# Is there any incremental benefit to conducting neuroimaging and neurocognitive assessments in the diagnosis of ADHD in young children? A machine learning investigation

**DOI:** 10.1101/2020.08.19.257600

**Authors:** Ilke Öztekin, Mark A. Finlayson, Paulo A. Graziano, Anthony S. Dick

## Abstract

Given the negative trajectories of early behavior problems associated with Attention-Deficit/Hyperactivity Disorder (ADHD), early diagnosis of ADHD is considered critical to enable early intervention and treatment. To this end, the current investigation employed machine learning to evaluate the relative predictive value of parent/teacher ratings, as well as behavioral and neural measures of executive function in predicting ADHD diagnostic category in a sample consisting of 162 young children (53.7% ADHD, ages 4 to 7, mean age 5.55, 67.9% male, 82.6% Hispanic/Latino). Among all the target measures assessed in the study, teacher ratings of executive function were identified as by far the most important measure in predicting ADHD diagnostic category. While a more extensive evaluation of neural measures, such as diffusion-weighted imaging, may provide more information as they relate to the underlying cognitive deficits associated with ADHD, the current study indicates that commonly used structural imaging measures of cortical thickness, as well as widely used cognitive measures of executive function, have little incremental value in differentiating typically developing children from those diagnosed with ADHD. Future research evaluating the importance of such measures in predicting children’s functional impairment in academic and social areas would provide additional insight into their contributing role in ADHD.

## INTRODUCTION

Attention-Deficit/Hyperactivity Disorder (ADHD) is a developmental disorder, characterized by a person’s difficulty to pay attention and control impulsive behavior (see Mahone & Denckla, 2017 for a historical overview). The National Resource Center on ADHD (NCR) currently lists fifteen million individuals as having ADHD in America. ADHD has a worldwide prevalence of 7.2% in children (Wolraich et al., 2019). Early identification and treatment of ADHD is considered critical due to the serious consequences of ADHD, including but not limited to academic/school and social difficulties (Hoza, 2007; Moffitt et al., 2002; Ros & Graziano, 2018; Shaw et al., 2003). Previous ADHD research implicates the important role of executive function (EF) processes (Barkley, 1997; Sergeant, 2000; Sonuga-Barke, 2002), with strong observed links between EF deficits and academic (Biederman et al., 2004; Blair & Razza, 2007; Clark et al., 2010) and social difficulties (Hill, 2004; Riggs et al., 2006). Machine learning techniques can leverage these important measures as a means to help identify 1) the most important predictors critical for successful classification of ADHD, and 2) provide objective measures of the relative clinical utility of target predictors. To this end, the current study evaluated the predictive utility of EF measures in three categories, namely parent/teacher ratings of EF, behavioral performance in EF tasks, and neural measures of cortical thickness in brain regions that support EF in a sample of 162 children between ages 4-7 (mean age 5.5 years).

An important advantage of multivariate modeling approaches in cognitive and translational neuroscience research has been their ability to combine information from data in a distributed fashion. As opposed to traditional univariate approaches, where data (such as signal intensity from functional MRI) are averaged over a specific region of interest (ROI), and compared to the mean signal in another ROI to assess potential differences in magnitude of neural activation, multivariate approaches use data from distributed patterns of a neural measure (e.g., brain activity or anatomical properties of the brain), and apply computational models to the data that extend beyond conducting statistical analyses on averages [for reviews see (Davis & Poldrack, 2013; Haxby, 2012; Haxby et al., 2001; Norman et al., 2006; Yang et al., 2012)]. While computationally more challenging, this latter approach enables researchers to answer key questions regarding mental representations (e.g., what type of information is represented in different brain regions and at different stages of cognitive processing) in cognitive neuroscience research (Haxby et al., 2001; Kuhl et al., 2011; Liang et al., 2013; Norman et al., 2006; Oztekin & Badre, 2011; Polyn et al., 2005)], and provides unique opportunities for clinical diagnosis and prediction of disorder or symptom severity, as well as behavioral outcomes associated with specific disorders. Despite this promising potential for such computational, multivariate approaches to advance translational neuroscience, significant challenges and pitfalls have prevented the development of generalizable methods and approaches that can be applied in clinical settings [see (Arbabshirani et al., 2017; Kassraian-Fard et al., 2016; Varoquaux, 2018; Varoquaux et al., 2017; Woo et al., 2017)].

One commonly recognized obstacle is *small sample size*, which leads to overfitting. Overfitting refers to the modeling of noise in the data rather than the underlying pattern of interest, which results in good performance on the training data, but poor performance on previously unseen data. Accordingly, overfitting is a major obstacle in generating a valid predictive model that can produce generalizable results for clinical diagnosis, and which can be used across a variety of clinical settings (Arbabshirani et al., 2017; Foster et al., 2014; Kassraian-Fard et al., 2016; Pulini et al., 2019; Varoquaux, 2018; Varoquaux et al., 2017; Woo et al., 2017). Indeed, most neuroscience studies suffer from small sample sizes (Arbabshirani et al., 2017; Foster et al., 2014; Kassraian-Fard et al., 2016; Pulini et al., 2019; Varoquaux, 2018). Notably, the current investigation leveraged a data set with a larger sample size (162 participants) that provided a unique opportunity to leverage predictive modeling that utilized a machine learning approach.

Another salient pitfall centers around the selection of features in the model. Two important factors stand out with respect to this obstacle (Arbabshirani et al., 2017; Foster et al., 2014; Woo et al., 2017). The first is *unbiased feature selection*. In their critical review of predictive modeling approaches for clinical diagnosis, Arbabshirani et al. (Arbabshirani et al., 2017) and Woo et al. (Woo et al., 2017) point out that it is common for predictive modeling studies to use a group analysis over the whole dataset to select appropriate features for the classifier. This, however, constitutes “double dipping” and produces biased results. In our approach, our feature selection was performed *a priori*, based on the ADHD literature that highlights the importance of executive function for ADHD, and prior research on the neurobiology of executive function. The second obstacle is *the need for theoretical motivation for selection of features*. In addition to selecting features in an unbiased manner, several critique articles (Foster et al., 2014; Kassraian-Fard et al., 2016; Woo et al., 2017) have put forth the importance of theoretical relevance of the features in the model, especially in the context of clinical diagnosis. Accordingly, in the current investigation, we limited our features to those that are theoretically relevant, namely the cognitive and neurobiological measures of executive function.

### Current Study

The ADHD Heterogeneity of Executive Function and Emotion Regulation Across Development (AHEAD) study aims to characterize the heterogeneity of well-established predictors of ADHD among young children (ages 4-to-7, mean age 5.5 years) across multiple levels of analysis. It thus uniquely provides multiple measures of executive function, which we focus in the current investigation, at both behavioral/cognitive and neural levels at an age-range where early diagnosis of ADHD is critically important. Notably, several research groups have previously applied machine learning to better characterize the heterogeneity of the behavioral profile in ADHD [for instance see (Karalunas et al., 2019; Qureshi et al., 2016)]. However, to date, none of the attempts have been adapted for clinical diagnosis in children with early diagnosis—i.e., in the pre-K age range during which initial contact with educators and clinicians typically occurs. Our AHEAD study provides a unique data set with the potential for a significant contribution in this domain. The AHEAD study is among the first to scan children with ADHD as young as 4–7 years. The age range in existing studies is usually 9 or older, with only a few studies sampling an age range of 7 and above (Fair et al., 2012; Karalunas et al., 2014; Karalunas et al., 2019; Qureshi et al., 2017). Given that early diagnosis of ADHD is critical, the current investigation provides a unique opportunity to assess whether predicting ADHD from scans is feasible in this age range.

#### Executive Function and ADHD

Importantly, previous research points to the critical role of executive function (EF) processes in ADHD (Barkley, 1997; Sergeant, 2000; Sonuga-Barke, 2002). Crucially, a strong relationship has been noted between EF deficits and academic (Biederman et al., 2004; Blair & Razza, 2007; Clark et al., 2010), as well as social difficulties (Hill, 2004; Riggs et al., 2006). With respect to the neurobiological markers that support EF, previous research notes the functional interactions among lateral frontal, inferior frontal/insular (Aron et al., 2004), medial frontal/anterior cingulate/pre-SMA (Aron et al., 2004; Bunge & Wright, 2007; Fedota et al., 2014; Miller & Cohen, 2001; Rushworth et al., 2005), lateral parietal (Corbetta & Shulman, 2002), and dorsal striatal (Morein-Zamir & Robbins, 2015) regions of the brain (Hart et al., 2014). Accordingly, our machine learning approach in this investigation exclusively focuses on the cognitive behavioral and neural measures of executive function, with the primary goal of assessing the utility of predicting ADHD diagnostic category from the target measures of executive function. Within this overarching goal, we pursued two aims: 1) to evaluate the predictive value of behavioral and neural correlates of executive function, and 2) to determine the relative importance of each measure in predicting ADHD diagnostic category. Given the implicated importance of executive function in ADHD, we hypothesized that our models can successfully predict ADHD diagnostic category.

## MATERIALS AND METHODS

### Participants and Recruitment

Children and their caregivers were recruited from local schools and mental health agencies via brochures, radio and newspaper ads, and open houses/parent workshops. Legal guardians contacted the clinic and were directed to the study staff for screening questions to determine eligibility. For the ADHD sample, if the parent (1) endorsed clinically significant levels of ADHD symptoms (six or more symptoms of either Inattention or Hyperactivity/Impulsivity according to the DSM-5 OR a previous diagnosis of ADHD), (2) indicated that the child is currently displaying clinically significant academic, behavioral, or social impairments as measured by a score of 3 or higher on a seven-point impairment rating scale (Fabiano et al., 2006), and (3) were not taking any psychotropic medication, the parent and child were invited to participate in an assessment to determine study eligibility. For the typically developing sample, if the parent (1) endorsed less than 4 ADHD symptoms (across either Inattention or Hyperactivity/Impulsivity according to the DSM-5), (2) less than 4 Oppositional Defiant Disorder (ODD) symptoms, and (3) indicated no clinically significant impairment (score below 3 on the impairment rating scale), the parent and child were invited to participate in an assessment to determine study eligibility. Participants were also required to be enrolled in school during the previous year, have an estimated IQ of 70 or higher, have no confirmed history of an Autism Spectrum Disorder, and be able to attend an 8-week summer treatment program prior to the start of the next school year (ADHD groups only).

During intake, ADHD diagnosis (and comorbid disruptive behavior disorders) was assessed through a combination of parent structured interview (Computerized-Diagnostic Interview Schedule for Children (Shaffer et al., 2000) and parent and teacher ratings of symptoms and impairment (Disruptive Behavior Disorders Rating Scale, Impairment Rating Scale (Fabiano et al., 2006) as is recommended practice (Pelham et al., 2005). Dual Ph.D. level clinician review was used to determine diagnosis and eligibility. The final sample included 162 children (87 with ADHD, and 75 healthy developing controls). Among the participants with ADHD, 61 (70%) had comorbid ODD diagnosis. Parental consent and assent was obtained in accordance with the Office of Research Integrity at Florida International University.

### Measures of Executive Function (EF)

#### 1. Parent/Teacher Ratings of EF

We utilized the Emergent Metacognition Index (MCI) *t*-score from the Behavior Rating Inventory of Executive Function (BRIEF, BRIEF-P, Gioia, Espy & Isquith, 2003) for our measure of parent and teacher ratings of executive function. The MCI is thought to reflect the ability to maintain information and/or activities in working memory, as well as to plan and organize problem-solving approaches. In the BRIEF Preschool, The MCI is composed of the Working Memory and Plan/Organize scales. In the BRIEF Child, the MCI is composed of the Initiate, Working Memory, Plan/Organize, Organization of Materials, and Monitor scales. In our sample, BRIEF-P was completed for 115 of our participants (ages 4-5; Chronbach’s alpha for MCI .976 for BRIEF-P Teacher and .970 for BRIEF-P Parent), and BRIEF Child was completed for 47 of our participants (ages 6-7; Chronbach’s alpha for MCI.724 for Teacher and .978 for Parent).

#### 2. Behavioral/Cognitive Measures of EF

Our dataset allowed us to evaluate performance in three tasks that measure executive function:

i. *The Flanker Task*. The Flanker Task of Inhibitory Control and Attention is part of the EF assessments utilized in the NIH Toolbox-Cognition battery (http://www.nihtoolbox.org). This task measures the participant’s responses under conditions where the surrounding stimuli and target are either congruent or incongruent.
ii. *The Dimensional Change Card Sorting Task (DCCS)*. The DCCS is also part of the Cognition battery of the NIH Toolbox. Participants sort objects by color or shape. Both the Flanker and DCCS tasks have been validated with children as young as age 3 (Zelazo et al., 2013).
iii. *Head-Toes-Knees-Shoulders (HTKS) Task*. HTKS (Cameron Ponitz et al., 2008) is a brief and widely-used task with young children ages 4-7 that assesses multiple aspects of EF. This task implements pairings of behavioral rules: “touch your head” and “touch your toes”, “touch your shoulders” and “touch your knees”. The children first respond naturally, and then are asked to switch the rules by responding in the opposite manner. Graziano and colleagues (Graziano et al., 2015) have previously established the ecological validity of this task in young children with ADHD.

#### 3. Neurobiological Measures

All imaging was performed using a research-dedicated 3 Tesla Siemens MAGNETOM Prisma MRI scanner (V11C) with a 32-channel coil located on the university campus. Children first completed a preparatory phase using realistic mock scanner in the room across the hall from the magnet. Here they were trained to stay still, and were also acclimated to the enclosed space of the magnet, the back projection visual presentation system, and to the scanner noises (in this case, presented with headphones). When they were properly trained and acclimated, they were moved to the magnet. In the magnet, during the structural scans, children watched a child-friendly movie of their choice. Ear protection was used, and sound was presented through MRI compatible headphones.

Structural MRI scans were collected using a 3D T1-weighted sequence (axial; 1 × 1 × 1 mm) with prospective motion correction (Siemens vNAV; Tisdall et al., 2012), according to the Adolescent Brain and Cognitive Development (ABCD) protocol (Hagler et al., 2019). Cortical thickness measures were extracted in target regions that support executive function. To provide a semi-automated parcellation of the cerebral cortices and volume of subcortical structures, we constructed two-dimensional surface renderings of each participant's brain using FreeSurfer (Dale et al., 1999; Fischl & Dale, 2000). We computed cortical thickness as part of the standard FreeSurfer reconstruction pipeline (Rohde et al., 2004), as this has been shown to have high correspondence to histological measurements of cortical thickness (Yeh et al., 2010).

### Quality Control of Magnetic Resonance Imaging Scans

Movement artifacts in T1-weighted MRI scans are common, especially in pediatric populations in this age range, and especially in children with ADHD. Fortunately, FreeSurfer is robust to movement-related artifacts, as, except in extreme cases, the program is able to accurately identify intensity differences between white matter and grey matter inherent in the T1-weighted image. In some cases, however, manual intervention is necessary. In this manual intervention, each individual MRI scan is inspected, and in cases where the program does not adequately identify the appropriate regional boundaries, manual edits are employed. We also visually rated each T1-weighted image on a seven-point scale ranging from “Poor = 1” to “Excellent = 4”, with allowances for half-points (e.g., 3.5). Scans for both groups were generally rated “Very Good” to “Excellent”, with an average of 3.61 for the ADHD group, and 3.54 for the typical control group. There were no significant group differences for the quality of the scans, *t*(145.46) = -0.72, *p*= 0.47.

### Machine Learning Approach

#### Features

Following the strong link between ADHD and executive function observed in prior research, our features constitute parent/teacher ratings of executive function (the Emergent Metacognition Composite score from BRIEF), performance in the three tasks that measure executive function (Flanker task, Dimensional Card Sorting task, HTKS task), and cortical thickness measures in the subregions of the frontal (superior frontal gyrus, middle frontal gyrus, inferior frontal gyrus-pars opercular, inferior frontal gyrus-pars triangularis, inferior frontal gyrus-pars orbitalis, insula, anterior cingulate, middle-anterior cingulate, middle-posterior cingulate, rectus) and parietal (superior parietal lobule, intraparietal and transverse parietal sulci, angular gyrus, supramarginal gyrus, and precuneus) cortices in both hemispheres. In addition to our target features, we also evaluated two demographic features, namely age and gender.

#### Model Construction and Assessment

We employed the *scikit-learn* (https://scikit-learn.org/stable/) open source machine learning platform for constructing our models. In order to evaluate feature importance from classifier coefficients, we adapted a Support Vector Machine classifier with a linear kernel. For model validation, we leveraged the built-in cross-validation function of the scikit-learn library. Performance was then evaluated with the commonly employed accuracy scores, as well as F1 scores obtained across the cross-validation indices for each model. F1 scores are a classification performance metric that is calculated based on the precision (p) and recall (r), 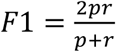 Following the recommended practice in the field, our primary assessment of statistical significance employed permutation tests (Combrisson & Jerbi, 2015; Noirhomme et al., 2014; Pereira et al., 2009). 5000 permutations were employed for each evaluated model. Along with permutation tests, we also report two baseline accuracy levels (see Table 1) as additional benchmark comparisons, namely the baseline for ADHD in our current sample, and the baseline for ADHD in the population. Our dataset consists of 87 participants with ADHD, and 75 typically developing (TD) children. Accordingly, the baseline for the present data set is .537. The population baseline, derived from the pooled worldwide prevalence of ADHD among children (Wolraich et al., 2019) is .072.

## RESULTS

Below we assess our target measures and their predictive utility for ADHD classification. Our analytic plan follows a two-step approach to determine model performance and relative predictive utility of our target features: *1) Feature Elimination*. To this end, we ran multiple models and compared the performance metrics across the models. This approach pursued four categories of models, i) a model including the demographics features (age and gender), ii) a model including the parent/teacher ratings of EF (Emergent Metacognition Composite t-score from BRIEF), iii) a model including the behavioral/cognitive measures of EF (including scores from HTKS, Flanker and Dimensional Card Sorting tasks), and iv) neural models assessing cortical thickness measures in our target regions for EF. *2) Feature Importance Rankings.* Within each model, we were able to further assess the relative importance of our features for predicting diagnostic category. We utilized the coefficients of our linear SVM classifier as an index for the relative importance across our features within each model.

**Table 1.** Model performance metrics across the four sets of target features.

| Models | Performance Metric |  |  |  | Model Evaluation |  |  |
| --- | --- | --- | --- | --- | --- | --- | --- |
|  | Acc | F<sub>1</sub> Macro | F<sub>1</sub> ADHD | F<sub>1</sub> TD | PT | CSB | PPB |
| 1. Demog. | .574 | .551 | .651 | .452 | $p < .047$ | .537 | .072 |
| 2. P/T Rat. | .926 | .925 | .929 | .921 | $p < .001$ | .537 | .072 |
| 3. EF Tasks | .667 | .666 | .674 | .658 | $p < .001$ | .537 | .072 |
| 4. Neural | .612 | .608 | .641 | .575 | $p < .016$ | .537 | .072 |

PT = Permutation Tests
CSB = Baseline for ADHD in the current study
PPB = Baseline for prevalence of ADHD in the population

### Predicting ADHD diagnostic category

For each model, a Support Vector Machine classifier with a linear kernel was trained to distinguish the two diagnostic categories (ADHD diagnosis absent, ADHD diagnosis present) using the target features. We scaled our features using the MinMaxScaler function built in the sci-kit machine learning library. To evaluate the performance of our models, we conducted cross validation across five indices (k-folds). Adapting five-folds follows the recommendation of Varoquax and colleagues (Varoquaux, 2018; Varoquaux et al., 2017), whose work suggested random splits with 20% data yield the best cross validation results in machine learning applications in neuroscience research.

To evaluate feature importance rankings, we leveraged the magnitude of classifier coefficients across the target features in the model. Weights obtained from the resultant classifier coefficients of linear SVM classifiers can be utilized to infer feature importance rakings within the assessed model. Thus, the average classifier coefficients across the five cross-validation indices provided an index for the feature importance rankings for each model evaluated. Figure 1 illustrates the feature importance rankings and classifier performance across the four sets of models, described in more detail below.

**Figure 1.**
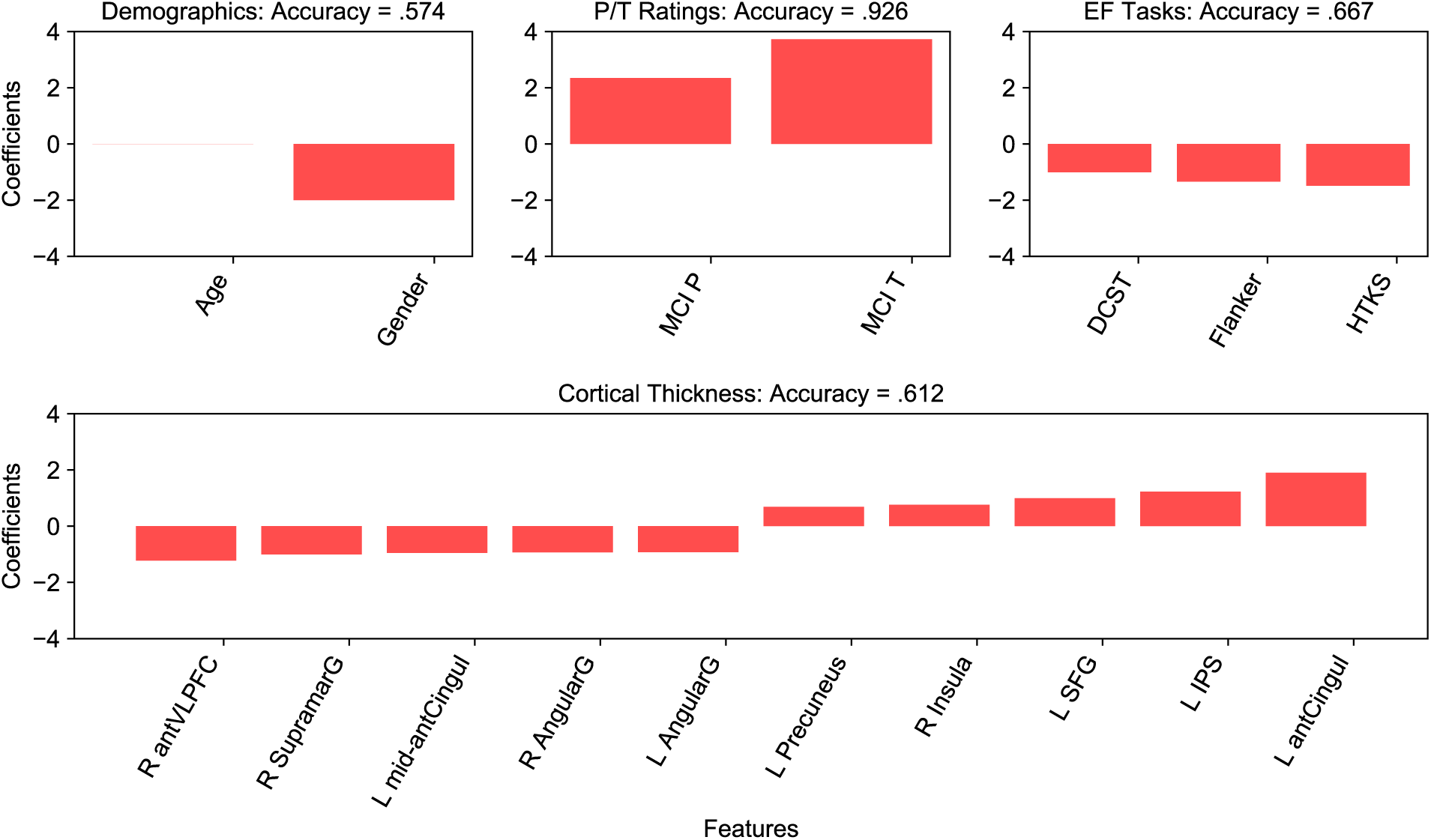
Accuracy and feature importance rankings from the four sets of models (1-demographics, 2-parent/teacher ratings of EF as measured by the Meta Cognition Index of the BRIEF assessment, 3-EF tasks, Dimensional Card Sorting. Flanker, and Head-Toes-Knees-Shoulders tasks, 4-Neural Measures, namely cortical thickness measures (the top 10 most important regions are shown).

#### Model 1: Demographics

The demographics model included two features, age and gender. Across the five cross-validation indices, this model yielded an average accuracy of .574 (*p* < .047), with gender yielding classifier coefficients of higher magnitude than age (see Figure 1).

#### Model 2: Parent and Teacher Ratings of Executive Function

Our second model included the Emergent Metacognition Index t-scores from the BRIEF parent and BRIEF teacher ratings. This model achieved an average accuracy score of .926 (*p* < .001), with the teacher ratings yielding higher classifier coefficients compared to parent ratings (Figure 1).

#### Model 3: Behavioral/Cognitive Measures of Executive Function

Our EF tasks included the DCCS, the Flanker task, and the Head-toes Knees-Shoulder Task (HTKS). This model leveraged the behavioral performance across the three tasks in order to distinguish ADHD diagnostic category. This model achieved an average accuracy score of .667 (*p* < .001), with the HTKS and Flanker tasks yielding classifier coefficients of greater magnitude than the DCCS task.

#### Model 4: Neural Measures of Cortical Thickness

Our next set of models explored our neural measures of cortical thickness in target regions that support executive function. We first ran a model including cortical thickness measures in both hemispheres of the brain. Figure 1 depicts classifier performance (average accuracy .612, *p* < .016), and coefficients for the top 10 features in this model. We next ran two models that contained features from the left and right hemispheres respectively. The model assessing cortical thickness measures from the left hemisphere yielded an average accuracy of .612 (*p* < .01), while the model evaluating cortical thickness measures from the right hemispheres achieved an accuracy of.50 (*p* > .64). Accordingly, our current investigation points to the more pronounced importance of the left hemisphere in predicting diagnostic category. In the left hemisphere, the regions yielding the top three classifier coefficients were the anterior cingulate, the intraparietal transverse sulcus, and the superior frontal gyrus (Figure 1).

#### Model 5: Full Model

Among the four sets of models evaluated, our analyses implicate the BRIEF Emergent Metacognition ratings (with teacher ratings ranking higher than the parent ratings in predicting diagnostic category) to be the most important sets of features. This model achieved .926 accuracy. We next evaluated whether the classifier’s performance could be further improved by including additional features from our target measures. This full model included the demographics, the parent/teacher ratings of MCI, cognitive measures of EF, and cortical thickness in the left anterior cingulate, the left intraparietal transverse sulcus and the left superior frontal gyrus. Across the five cross-validation indices, this full model achieved an average accuracy of .944 (p < .001). Table 1 summarizes the performance metrics across the models.

In summary, our computational modeling approach implicates the Emergent Metacognition scores from the BRIEF assessment to be the most important target measure. More specifically, the teacher ratings were identified to be the most important feature. In our current set of findings, the two BRIEF ratings alone achieved an accuracy of .926. Adding additional features that included demographics, performance in EF tasks and neural measures of cortical thickness in regions identified to be important in predicting ADHD diagnostic category increased the classifier’s performance to .944. Thus, when considering different types of variables (e.g., demographics, parent/teacher ratings of EF, behavioral measures of EF, and neural measures of cortical thickness in regions that support EF) and their relative importance for classification of diagnostic category, our current set of analyses point to the critical importance of teacher ratings of executive function for classification of ADHD diagnostic category.

### High comorbid ODD in the ADHD sample

Recall that our ADHD sample has high (70%) comorbid ODD. As such, one concern that could potentially limit the implications of our results is that the high prevalence of ODD might have contributed to the differentiation of diagnostic category in our models. Unfortunately, due to the low sample size of participants with only ADHD diagnosis (n = 26), it was not methodologically feasible/sound to run our models to distinguish the 3 diagnostic categories (i.e. 1-ADHD diagnosis absent, 2-only ADHD diagnosis present, and 3-ADHD + ODD diagnosis present). To assess the potential impact/change in the performance of our models reported above, we repeated our analyses by excluding the 26 participants with only ADHD diagnosis. Thus, we trained our classifiers to distinguish ADHD diagnostic category across our typically developing sample and children with ADHD+ODD diagnosis. This set of models yielded similar results [accuracy = .918, p < .001 for parent/teacher ratings of EF, accuracy = .711, p < .001 for cognitive EF tasks, accuracy = .607, p < .045 for the neural measures of cortical thickness], except for the demographics model [accuracy = .511 p > .85] which was no longer significant.

We once again acknowledge that the most ideal approach to eliminate this potential confound would be to run models trained to distinguish the three diagnostic categories. Unfortunately, in our current data set, this was not possible due to the low number of participants in the ADHD-only category. Future work utilizing larger samples will be fundamental in thoroughly addressing comorbid ODD within the ADHD sample.

## DISCUSSION

Given the negative trajectories of early behavior problems associated with ADHD, as well as its high public health cost, early diagnosis of ADHD is critical to enable early intervention and treatment. The current investigation aimed to evaluate the feasibility of predictive modeling using cognitive, behavioral and neurobiological measures of executive function for categorical classification of ADHD in young children. Notably, the current set of findings are the first to employ this approach in Pre-K children. The AHEAD study is among the first to scan children with ADHD as young as 4–7 years (mean age 5.5 years) with a good sample size (n > 160). Given that early diagnosis of ADHD is critical, the current study provides a unique opportunity to assess whether predicting ADHD from scans is feasible in this age range.

Results from our machine learning models indicated that the full model including features from all sets of target features yielded a .944 performance in predicting diagnostic category. Crucially, most of this high performance is accounted by the parent/teacher BRIEF Emergent Metacognition Index scores, which alone achieved a performance of .926. These findings implicate the critical importance of this measure in predicting diagnostic category, and this has implications for clinical diagnosis. Thus, the current set of results suggest that the rating scales are enough to distinguish the presence or absence of ADHD, and that more expensive and extensive behavioral and neural testing might not be necessary. In particular, our models identified the teacher ratings to be most diagnostic in predicting diagnostic category, further suggesting that the functional impairments pertaining to executive function processes experienced in school can be clearly differentiated among children with ADHD from their typically developing peers.

From one perspective, it is not surprising that parent-teacher ratings on one assessment (i.e., BRIEF) are good at predicting diagnostic category that is itself predicated, in part, on parent-teacher ratings on another assessment (e,g., DBD rating scale). Thus, for simple categorical diagnosis, collection of additional measures does not provide substantial improvement in the detection of ADHD. This is important because this study addresses some of the critical questions raised by the American Academy of Pediatrics Subcommittee on Children and Adolescents with ADHD. In their recent Clinical Practice Guideline (Wolraich et al., 2019, p. 3) asked 1) what is the comparative diagnostic accuracy of approaches that can be used in the primary care practice setting or by specialists to diagnose ADHD among children younger than 7 years of age?; 2) what is the comparative diagnostic accuracy of EEG, imaging, or executive function approaches that can be used in the primary care practice setting or by specialists to diagnose ADHD among individuals aged 7 to their 18th birthday?; 3) are there more formal neuropsychological, imaging, or genetic tests that improve the diagnostic process? (Wolraich et al., 2019). Our results map on to the committee’s recommendations for young children in terms of the reliance on parent and teacher ratings scales for aiding clinicians in the diagnosis of ADHD. Thus, while data from our study shows that neurocognitive and imaging measures are not incrementally useful for simply detecting the presence or absence of ADHD, more research is needed to determine their utility as it relates to predicting more functional outcomes (e.g., academic/social impairments) as well as comorbid conditions.

More specifically, the classifier coefficients in our models allowed us to further rank our target features in their relative importance for predicting ADHD diagnostic category. Among the executive functions measured, performance in the HTKS task yielded the highest coefficient in our current investigation, and the Flanker task was identified as more diagnostic than the DCCS. With respect to the neural measures, regions of the left hemisphere were more diagnostic compared to the regions in the right hemisphere of the brain. Among our target regions important for executive function, the top regions identified were the left anterior cingulate, the left superior frontal gyrus and the left intraparietal transverse sulcus.

Subparts of the cingulate cortex, including the anterior and middle-posterior cingulate cortex have been previously implicated in ADHD research (Rubia, 2018). This region has been previously associated with cognitive and impulse control, and a decreased connectivity between the anterior cingulate and the posterior parietal cortex in ADHD has been observed in prior research [see Vogt (2019) for a review]. Given its implication in executive function and previous ADHD research, it is not surprising that our classifier indicated this region as one of the top predictors among the cortical thickness measures evaluated in our study.

The intraparietal sulcus is a very significant contributor to working memory (Chein et al., 2003; Cowan, 2016; Cowan et al., 2011; Oztekin et al., 2009; Smith & Jonides, 1998; Xu & Chun, 2006), and working memory deficits have been commonly observed in previous ADHD research (Hammer et al., 2015; Karalunas et al., 2017; Palladino & Ferrari, 2013; Raiker et al., 2019; Raiker et al., 2012). Notably, this region has been previously implicated for its importance in supporting focus of attention during WM operations in healthy adults (Cowan et al., 2011; Oztekin et al., 2009). Given the widely-established deficits in working memory with ADHD, future research would benefit from a deeper investigation of the specific role of this region in potentially modulating ADHD related effects in working memory function.

The superior frontal gyrus (SFG), located at the superior part of the prefrontal cortex is anteromedially connected to the anterior and mid-cingulate. The dorsolateral portion of the SFG is connected with the dorsolateral and ventrolateral prefrontal cortex, and the posterior portion of the SFG is part of the motor control network (Li et al., 2013). Therefore, it has connections to significant regions that modulate motor, cognitive control and executive function. In addition, it is part of a relatively recently identified fiber pathway, the frontal aslant tract (FAT). Recent work from our group (Dick et al., 2019; Garic et al., 2019) has implicated FAT to be an important potential moderator for ADHD related deficits in executive function. Our modeling results implicate SFG as a potential diagnostic region. Unfortunately, our current parcellation did not allow us to independently evaluate the subregion of SFG. We thus note that future research would benefit from a more focused assessment of this region.

### Directions for Future Research

The current set of findings were a first in exploring the utility of predictive modeling for ADHD classification in the young age-range of 4-7 years. As it relates to neuroimaging, we explored cortical thickness measures which provided slightly more predictive power towards determining the presence of ADHD as simply knowing a child’s age and sex (.612 versus .574 accuracy). Thus, we acknowledge the lack of clinical utility of cortical thickness measures. Whether the inclusion of more structural, and diffusion-weighted-imaging (DWI) measures yields better clinical utility is a question for further research to examine. However, to date, no predictive modeling approach has evaluated DWI measures for classification of ADHD. Notably, most computational modeling and machine learning approaches in the existing literature have focused on functional MRI data (most commonly functional connectivity analysis of resting state data). Accordingly, a major criticism has been the issue of transportability (Foster et al., 2014; Woo et al., 2017), and that the findings do not have the potential to generalize to or be easily applied in clinical settings. As such, the inclusion of DWI measures could potentially fill this important gap in the literature.

Acknowledging that neuroimaging may yield only minimal clinical utility, there are still reasons to continue to pursue research within this domain. First, a comprehensive modeling approach should be able to identify which neural measures are more diagnostic, and how they compare to non-neural measures, such as behavioral assessments of theoretically important constructs for ADHD, such as executive function. Second, the ability to derive the most diagnostic neural measures for ADHD classification can facilitate researchers to further investigate the direct link between these regions and the specificity of their functional outcomes are with respect to ADHD-related impacts on the corresponding cognitive constructs. Complementary efforts that utilize and combine predictive modeling approaches with further assessments of the implicated regions/neural measures in how they modulate ADHD-related functional outcomes can facilitate unique ways to define and understand this disorder, and may foster development of novel approaches to the classification and treatment of ADHD.

### Conclusion

The AHEAD study is among the first to provide executive function measures for young children with ADHD (ages 4-7) at multiple levels of analysis. Providing a large sample size with complementary measures for both cognitive/behavioral and neural measures of executive function, the AHEAD study enabled the first assessment of the feasibility of predictive modeling and machine learning approaches in diagnosis of ADHD in Pre-K children. Our results suggest the critical importance of teacher ratings of executive function in predicting ADHD diagnostic category. Our models further allowed ranking of the importance of our target behavioral and neural measures of executive function. Among the three executive function tasks evaluated, feature importance rankings implicated performance in the HTKS to be the most diagnostic executive function measure. With respect to neural measures, our analyses focused on cortical thickness measures as the target neurobiological assessment. While a more extensive evaluation of neural measures, such as diffusion-weighted imaging, may provide more information as it relates to the underlying cognitive deficits associated with ADHD, the current study indicates that commonly used cortical thickness measures of structural imaging, as well as commonly employed cognitive tasks measuring EF, might have little incremental value in differentiating typically developing children from those diagnosed with ADHD. While teacher ratings of EF are more cost effective and provided a strong prediction of diagnostic status, it is important to acknowledge some of the measurement overlap in the EF items assessed in the BRIEF questionnaire and those present in the DSM-5 criteria for ADHD. Thus, it will be important for future work to not only focus on the utility of neural, neurocognitive, and behavioral measures of EF as it relates to diagnostic classification, but most importantly how they may predict children’s functional impairment in areas such as academic and social functioning.

## ACKNOWLEDGEMENTS

The AHEAD study is funded by grants from the National Institute of Health (R56MH108616, R01MH112588 and R01DK119814) to P. A. Graziano and A. S. Dick. This research was supported by a Computational Administrative Supplement to parent grant R01MH112588.

## REFERENCES

Arbabshirani, M. R., Plis, S., Sui, J., & Calhoun, V. D. (2017). Single subject prediction of brain disorders in neuroimaging: Promises and pitfalls. Neuroimage, 145(Pt B), 137–165. https://doi.org/10.1016/j.neuroimage.2016.02.079

Aron, A. R., Robbins, T. W., & Poldrack, R. A. (2004). Inhibition and the right inferior frontal cortex. Trends Cogn Sci, 8(4), 170–177. https://doi.org/10.1016/j.tics.2004.02.010

Barkley, R. A. (1997). Behavioral inhibition, sustained attention, and executive functions: constructing a unifying theory of ADHD. Psychol Bull, 121(1), 65–94. https://doi.org/10.1037/0033-2909.121.1.65

Biederman, J., Monuteaux, M. C., Doyle, A. E., Seidman, L. J., Wilens, T. E., Ferrero, F., Morgan, C. L., & Faraone, S. V. (2004). Impact of executive function deficits and attention-deficit/hyperactivity disorder (ADHD) on academic outcomes in children. J Consult Clin Psychol, 72(5), 757–766. https://doi.org/10.1037/0022-006X.72.5.757

Blair, C., & Razza, R. P. (2007). Relating effortful control, executive function, and false belief understanding to emerging math and literacy ability in kindergarten. Child Dev, 78(2), 647–663. https://doi.org/10.1111/j.1467-8624.2007.01019.x

Bunge, S. A., & Wright, S. B. (2007). Neurodevelopmental changes in working memory and cognitive control. Curr Opin Neurobiol, 17(2), 243–250. https://doi.org/10.1016/j.conb.2007.02.005

Cameron Ponitz, C. E., McClelland, M. M., Jewkes, A. M., Connor, C. M., Farris, C. L., & Morrison, F. J. (2008). Touch your toes! Developing a direct measure of behavioral regulation in early childhood. Early Childhood Research Quarterly, 23(2), 141–158. https://doi.org/https://doi.org/10.1016/j.ecresq.2007.01.004

Chein, J. M., Ravizza, S. M., & Fiez, J. A. (2003). Using neuroimaging to evaluate models of working memory and their implications for language processing. Journal of Neurolinguistics, 16(4–5), 315–339. https://doi.org/10.1016/S0911-6044(03)00021-6

Clark, C. A. C., Pritchard, V. E., & Woodward, L. J. (2010). Preschool executive functioning abilities predict early mathematics achievement. Dev Psychol, 46(5), 1176–1191. https://doi.org/10.1037/a0019672

Combrisson, E., & Jerbi, K. (2015). Exceeding chance level by chance: The caveat of theoretical chance levels in brain signal classification and statistical assessment of decoding accuracy. Journal of Neuroscience Methods, 250, 126–136. https://doi.org/https://doi.org/10.1016/j.jneumeth.2015.01.010

Corbetta, M., & Shulman, G. L. (2002). Control of goal-directed and stimulus-driven attention in the brain. Nat Rev Neurosci, 3(3), 201–215. https://doi.org/10.1038/nrn755

Cowan, N. (2016). Working Memory Maturation: Can We Get at the Essence of Cognitive Growth? Perspect Psychol Sci, 11(2), 239–264. https://doi.org/10.1177/1745691615621279

Cowan, N., Li, D., Moffitt, A., Becker, T. M., Martin, E. A., Saults, J. S., & Christ, S. E. (2011). A neural region of abstract working memory. J Cogn Neurosci, 23(10), 2852–2863. https://doi.org/10.1162/jocn.2011.21625

Dale, A. M., Fischl, B., & Sereno, M. I. (1999). Cortical surface-based analysis. I. Segmentation and surface reconstruction. Neuroimage, 9(2), 179–194. https://doi.org/10.1006/nimg.1998.0395

Davis, T., & Poldrack, R. A. (2013). Measuring neural representations with fMRI: practices and pitfalls. Ann N Y Acad Sci, 1296, 108–134. https://doi.org/10.1111/nyas.12156

Dick, A. S., Garic, D., Graziano, P., & Tremblay, P. (2019). The frontal aslant tract (FAT) and its role in speech, language and executive function. Cortex, 111, 148–163. https://doi.org/10.1016/j.cortex.2018.10.015

Fabiano, G. A., Pelham, J., William E, Waschbusch, D. A., Gnagy, E. M., Lahey, B. B., Chronis, A. M., Onyango, A. N., Kipp, H., Lopez-Williams, A., & Burrows-MacLean, L. (2006). A practical measure of impairment: Psychometric properties of the impairment rating scale in samples of children with attention deficit hyperactivity disorder and two school-based samples. Journal of Clinical Child and Adolescent Psychology, 35(3), 369–385.

Fair, D. A., Bathula, D., Nikolas, M. A., & Nigg, J. T. (2012). Distinct neuropsychological subgroups in typically developing youth inform heterogeneity in children with ADHD. Proc Natl Acad Sci U S A, 109(17), 6769–6774. https://doi.org/10.1073/pnas.1115365109

Fedota, J. R., Hardee, J. E., Perez-Edgar, K., & Thompson, J. C. (2014). Representation of response alternatives in human presupplementary motor area: multi-voxel pattern analysis in a go/no-go task. Neuropsychologia, 56, 110–118. https://doi.org/10.1016/j.neuropsychologia.2013.12.022

Fischl, B., & Dale, A. M. (2000). Measuring the thickness of the human cerebral cortex from magnetic resonance images. Proc Natl Acad Sci U S A, 97(20), 11050–11055. https://doi.org/10.1073/pnas.200033797

Foster, K. R., Koprowski, R., & Skufca, J. D. (2014). Machine learning, medical diagnosis, and biomedical engineering research - commentary. Biomed Eng Online, 13, 94. https://doi.org/10.1186/1475-925X-13-94

Garic, D., Broce, I., Graziano, P., Mattfeld, A., & Dick, A. S. (2019). Laterality of the frontal aslant tract (FAT) explains externalizing behaviors through its association with executive function. Dev Sci, 22(2), e12744. https://doi.org/10.1111/desc.12744

Graziano, P., Garb, L., Ros-Demarize, R., Hart, K., & Garcia, A. (2015). Executive Functioning and School Readiness Among Preschoolers With Externalizing Problems: The Moderating Role of the Student–Teacher Relationship. Early Education and Development, 27, 1–17. https://doi.org/10.1080/10409289.2016.1102019

Hagler Jr, D. J., Hatton, S., Cornejo, M. D. et al. (2019). Image processing and analysis methods for the Adolescent Brain Cognitive Development Study. NeuroImage, 202, 116091. https://doi.org/10.1016/j.neuroimage.2019.116091

Hammer, R., Cooke, G. E., Stein, M. A., & Booth, J. R. (2015). Functional neuroimaging of visuospatial working memory tasks enables accurate detection of attention deficit and hyperactivity disorder. Neuroimage Clin, 9, 244–252. https://doi.org/10.1016/j.nicl.2015.08.015

Hart, H., Chantiluke, K., Cubillo, A. I., Smith, A. B., Simmons, A., Brammer, M. J., Marquand, A. F., & Rubia, K. (2014). Pattern classification of response inhibition in ADHD: toward the development of neurobiological markers for ADHD. Hum Brain Mapp, 35(7), 3083–3094. https://doi.org/10.1002/hbm.22386

Haxby, J. V. (2012). Multivariate pattern analysis of fMRI: the early beginnings. Neuroimage, 62(2), 852–855. https://doi.org/10.1016/j.neuroimage.2012.03.016

Haxby, J. V., Gobbini, M. I., Furey, M. L., Ishai, A., Schouten, J. L., & Pietrini, P. (2001). Distributed and overlapping representations of faces and objects in ventral temporal cortex. Science, 293(5539), 2425–2430. https://doi.org/10.1126/science.1063736

Hill, E. L. (2004). Executive dysfunction in autism. Trends Cogn Sci, 8(1), 26–32.

Hoza, B. (2007). Peer functioning in children with ADHD. Ambul Pediatr, 7(1 Suppl), 101–106. https://doi.org/10.1016/j.ambp.2006.04.011

Karalunas, S. L., Fair, D., Musser, E. D., Aykes, K., Iyer, S. P., & Nigg, J. T. (2014). Subtyping attention-deficit/hyperactivity disorder using temperament dimensions: toward biologically based nosologic criteria. JAMA Psychiatry, 71(9), 1015–1024. https://doi.org/10.1001/jamapsychiatry.2014.763

Karalunas, S. L., Gustafsson, H. C., Dieckmann, N. F., Tipsord, J., Mitchell, S. H., & Nigg, J. T. (2017). Heterogeneity in development of aspects of working memory predicts longitudinal attention deficit hyperactivity disorder symptom change. Journal of Abnormal Psychology, 126(6), 774–792. https://doi.org/10.1037/abn0000292

Karalunas, S. L., Gustafsson, H. C., Fair, D., Musser, E. D., & Nigg, J. T. (2019). Do we need an irritable subtype of ADHD? Replication and extension of a promising temperament profile approach to ADHD subtyping. Psychological Assessment, 31(2), 236–247. https://doi.org/10.1037/pas0000664

Kassraian-Fard, P., Matthis, C., Balsters, J. H., Maathuis, M. H., & Wenderoth, N. (2016). Promises, Pitfalls, and Basic Guidelines for Applying Machine Learning Classifiers to Psychiatric Imaging Data, with Autism as an Example. Front Psychiatry, 7, 177. https://doi.org/10.3389/fpsyt.2016.00177

Kuhl, B. A., Rissman, J., Chun, M. M., & Wagner, A. D. (2011). Fidelity of neural reactivation reveals competition between memories. Proc Natl Acad Sci U S A, 108(14), 5903–5908. https://doi.org/10.1073/pnas.1016939108

Lee, S. S., Humphreys, K. L., Flory, K., Liu, R., & Glass, K. (2011). Prospective association of childhood attention-deficit/hyperactivity disorder (ADHD) and substance use and abuse/dependence: a meta-analytic review. Clin Psychol Rev, 31(3), 328–341. https://doi.org/10.1016/j.cpr.2011.01.006

Li, W., Qin, W., Liu, H., Fan, L., Wang, J., Jiang, T., & Yu, C. (2013). Subregions of the human superior frontal gyrus and their connections. NeuroImage, 78, 46–58. https://doi.org/10.1016/j.neuroimage.2013.04.011

Liang, M., Mouraux, A., Hu, L., & Iannetti, G. D. (2013). Primary sensory cortices contain distinguishable spatial patterns of activity for each sense. Nat Commun, 4, 1979. https://doi.org/10.1038/ncomms2979

Mahone, E. M., & Denckla, M. B. (2017). Attention-Deficit/Hyperactivity Disorder: A Historical Neuropsychological Perspective. Journal of the International Neuropsychological Society: JINS, 23(9–10), 916–929. https://doi.org/10.1017/S1355617717000807

Miller, E. K., & Cohen, J. D. (2001). An integrative theory of prefrontal cortex function. Annu Rev Neurosci, 24, 167–202. https://doi.org/10.1146/annurev.neuro.24.1.167

Moffitt, T. E., Caspi, A., Harrington, H., & Milne, B. J. (2002). Males on the life-course-persistent and adolescence-limited antisocial pathways: follow-up at age 26 years. Dev Psychopathol, 14(1), 179–207.

Morein-Zamir, S., & Robbins, T. W. (2015). Fronto-striatal circuits in response-inhibition: Relevance to addiction. Brain Res, 1628(Pt A), 117–129. https://doi.org/10.1016/j.brainres.2014.09.012

Noirhomme, Q., Lesenfants, D., Gomez, F., Soddu, A., Schrouff, J., Garraux, G., Luxen, A., Phillips, C., & Laureys, S. (2014). Biased binomial assessment of cross-validated estimation of classification accuracies illustrated in diagnosis predictions. NeuroImage: Clinical, 4, 687–694. https://doi.org/https://doi.org/10.1016/j.nicl.2014.04.004

Norman, K. A., Polyn, S. M., Detre, G. J., & Haxby, J. V. (2006). Beyond mind-reading: multi-voxel pattern analysis of fMRI data. Trends Cogn Sci, 10(9), 424–430. https://doi.org/10.1016/j.tics.2006.07.005

Oztekin, I., & Badre, D. (2011). Distributed Patterns of Brain Activity that Lead to Forgetting. Front Hum Neurosci, 5, 86. https://doi.org/10.3389/fnhum.2011.00086

Oztekin, I., McElree, B., Staresina, B. P., & Davachi, L. (2009). Working memory retrieval: contributions of the left prefrontal cortex, the left posterior parietal cortex, and the hippocampus. J Cogn Neurosci, 21(3), 581–593. https://doi.org/10.1162/jocn.2008.21016

Palladino, P., & Ferrari, M. (2013). Interference control in working memory: comparing groups of children with atypical development. Child Neuropsychol, 19(1), 37–54. https://doi.org/10.1080/09297049.2011.633505

Pelham, J., William E, Fabiano, G. A., & Massetti, G. M. (2005). Evidence-based assessment of attention deficit hyperactivity disorder in children and adolescents. Journal of Clinical Child and Adolescent Psychology, 34(3), 449–476.

Pereira, F., Mitchell, T., & Botvinick, M. (2009). Machine learning classifiers and fMRI: A tutorial overview. NeuroImage, 45(1, Supplement 1), S199–S209. https://doi.org/https://doi.org/10.1016/j.neuroimage.2008.11.007

Polyn, S. M., Natu, V. S., Cohen, J. D., & Norman, K. A. (2005). Category-specific cortical activity precedes retrieval during memory search. Science, 310(5756), 1963–1966. https://doi.org/10.1126/science.1117645

Pulini, A. A., Kerr, W. T., Loo, S. K., & Lenartowicz, A. (2019). Classification Accuracy of Neuroimaging Biomarkers in Attention-Deficit/Hyperactivity Disorder: Effects of Sample Size and Circular Analysis. Biol Psychiatry Cogn Neurosci Neuroimaging, 4(2), 108–120. https://doi.org/10.1016/j.bpsc.2018.06.003

Qureshi, M. N., Min, B., Jo, H. J., & Lee, B. (2016). Multiclass Classification for the Differential Diagnosis on the ADHD Subtypes Using Recursive Feature Elimination and Hierarchical Extreme Learning Machine: Structural MRI Study. PLoS One, 11(8), e0160697. https://doi.org/10.1371/journal.pone.0160697

Qureshi, M. N. I., Oh, J., Min, B., Jo, H. J., & Lee, B. (2017). Multi-modal, Multi-measure, and Multi-class Discrimination of ADHD with Hierarchical Feature Extraction and Extreme Learning Machine Using Structural and Functional Brain MRI. Front Hum Neurosci, 11, 157. https://doi.org/10.3389/fnhum.2017.00157

Raiker, J. S., Friedman, L. M., Orban, S. A., Kofler, M. J., Sarver, D. E., & Rapport, M. D. (2019). Phonological Working Memory Deficits in ADHD Revisited: The Role of Lower Level Information-Processing Deficits in Impaired Working Memory Performance. J Atten Disord, 23(6), 570–583. https://doi.org/10.1177/1087054716686182

Raiker, J. S., Rapport, M. D., Kofler, M. J., & Sarver, D. E. (2012). Objectively-measured impulsivity and attention-deficit/hyperactivity disorder (ADHD): testing competing predictions from the working memory and behavioral inhibition models of ADHD. J Abnorm Child Psychol, 40(5), 699–713. https://doi.org/10.1007/s10802-011-9607-2

Riggs, N. R., Jahromi, L. B., Razza, R. P., Dillworth-Bart, J. E., & Mueller, U. (2006). Executive function and the promotion of social-emotional competence. Journal of Applied Developmental Psychology, 27(4), 300–309. https://doi.org/http://dx.doi.org/10.1016/j.appdev.2006.04.002

Rohde, G. K., Barnett, A. S., Basser, P. J., Marenco, S., & Pierpaoli, C. (2004). Comprehensive approach for correction of motion and distortion in diffusion-weighted MRI. Magn Reson Med, 51(1), 103–114. https://doi.org/10.1002/mrm.10677

Ros, R., & Graziano, P. A. (2018). Social functioning in children with or at risk for attention deficit/hyperactivity disorder: A meta-analytic review. Journal of Clinical Child & Adolescent Psychology, 47(2), 213–235. https://doi.org/10.1080/15374416.2016.1266644

Rubia, K. (2018). Cognitive Neuroscience of Attention Deficit Hyperactivity Disorder (ADHD) and Its Clinical Translation [10.3389/fnhum.2018.00100]. Frontiers in Human Neuroscience, 12, 100.

Rushworth, M. F., Buckley, M. J., Gough, P. M., Alexander, I. H., Kyriazis, D., McDonald, K. R., & Passingham, R. E. (2005). Attentional selection and action selection in the ventral and orbital prefrontal cortex. J Neurosci, 25(50), 11628–11636. https://doi.org/10.1523/JNEUROSCI.2765-05.2005

Sergeant, J. (2000). The cognitive-energetic model: an empirical approach to attention-deficit hyperactivity disorder. Neurosci Biobehav Rev, 24(1), 7–12.

Shaffer, D., Fisher, P., Lucas, C. P., Dulcan, M. K., & Schwab-Stone, M. E. (2000). NIMH Diagnostic Interview Schedule for Children Version IV (NIMH DISC-IV): description, differences from previous versions, and reliability of some common diagnoses. J Am Acad Child Adolesc Psychiatry, 39(1), 28–38. https://doi.org/10.1097/00004583-200001000-00014

Shaw, D. S., Gilliom, M., Ingoldsby, E. M., & Nagin, D. S. (2003). Trajectories leading to school-age conduct problems. Dev Psychol, 39(2), 189–200.

Smith, E. E., & Jonides, J. (1998). Neuroimaging analyses of human working memory. Proc Natl Acad Sci U S A, 95(20), 12061–12068. https://doi.org/10.1073/pnas.95.20.12061

Sonuga-Barke, E. J. (2002). Psychological heterogeneity in AD/HD--a dual pathway model of behaviour and cognition. Behav Brain Res, 130(1–2), 29–36.

Tisdall, M. D., Hess, A. T., Reuter, M., Meintjes, E. M., Fischl, B., & van der Kouwe, A.J. (2012). Volumetric navigators for prospective motion correction and selective reacquisition in neuroanatomical MRI. Magnetic resonance in medicine, 68(2), 389–399. https://doi.org/10.1002/mrm.23228

Varoquaux, G. (2018). Cross-validation failure: Small sample sizes lead to large error bars. Neuroimage, 180(Pt A), 68–77. https://doi.org/10.1016/j.neuroimage.2017.06.061

Varoquaux, G., Raamana, P. R., Engemann, D. A., Hoyos-Idrobo, A., Schwartz, Y., & Thirion, B. (2017). Assessing and tuning brain decoders: Cross-validation, caveats, and guidelines. Neuroimage, 145(Pt B), 166–179. https://doi.org/10.1016/j.neuroimage.2016.10.038

Vogt, B. A. (2019). Chapter 16 - Cingulate impairments in ADHD: Comorbidities, connections, and treatment. In B. A. Vogt (Ed.), Handbook of Clinical Neurology (Vol. 166, pp. 297–314). Elsevier. https://doi.org/https://doi.org/10.1016/B978-0-444-64196-0.00016-9

Wolraich, M. L., Hagan, J. F., Allan C et al. (2019). Clinical practice guideline for the diagnosis, evaluation, and treatment of Attention-Deficit/Hyperactivity Disorder in children and adolescents. Pediatrics, 144(4), e20192528. https://doi.org/10.1542/peds.2019-2528

Woo, C. W., Chang, L. J., Lindquist, M. A., & Wager, T. D. (2017). Building better biomarkers: brain models in translational neuroimaging. Nat Neurosci, 20(3), 365–377. https://doi.org/10.1038/nn.4478

Xu, Y., & Chun, M. M. (2006). Dissociable neural mechanisms supporting visual short-term memory for objects. Nature, 440(7080), 91–95. https://doi.org/10.1038/nature04262

Yang, Z., Fang, F., & Weng, X. (2012). Recent developments in multivariate pattern analysis for functional MRI. Neurosci Bull, 28(4), 399–408. https://doi.org/10.1007/s12264-012-1253-3

Yeh, F. C., Wedeen, V. J., & Tseng, W. Y. (2010). Generalized q-sampling imaging. IEEE Trans Med Imaging, 29(9), 1626–1635. https://doi.org/10.1109/TMI.2010.2045126

Zelazo, P. D., Anderson, J. E., Richler, J., Wallner-Allen, K., Beaumont, J. L., & Weintraub, S. (2013). II. NIH TOOLBOX COGNITION BATTERY (CB): MEASURING EXECUTIVE FUNCTION AND ATTENTION. Monographs of the Society for Research in Child Development, 78(4), 16–33. https://doi.org/10.1111/mono.12032

